# Longitudinal deformation based morphometry pipeline to study neuroanatomical differences in structural MRI based on SyN unbiased templates

**DOI:** 10.1101/2024.08.12.607581

**Authors:** Jürgen Germann, Flavia Venetucci Gouveia, M. Mallar Chakravarty, Gabriel A. Devenyi

## Abstract

Morphometric measures in humans derived from magnetic resonance imaging (MRI) have provided important insights into brain differences and changes associated with development and disease in vivo. Deformation-based morphometry (DBM) is a registration-based technique that has been shown to be useful in detecting local volume differences and longitudinal brain changes while not requiring a priori segmentation or tissue classification. Typically, DBM measures are derived from registration to common template brain space (one-level DBM). Here, we present a two-level DBM technique: first, the Jacobian determinants are calculated for each individual input MRI at the subject level to capture longitudinal individual brain changes; then, in a second step, an unbiased common group space is created, and the Jacobians co-registered to enable the comparison of individual morphological changes across subjects or groups. This two-level DBM is particularly suitable for capturing longitudinal intra-individual changes in vivo, as calculating the Jacobians within-subject space leads to superior accuracy. Using artificially induced volume differences, we demonstrate that this two-level DBM pipeline is 4.5x more sensitive in detecting longitudinal within-subject volume changes compared to a typical one-level DBM approach. It also captures the magnitude of the induced volume change much more accurately. Using 150 subjects from the OASIS-2 dataset, we demonstrate that the two-level DBM is superior in capturing cortical volume changes associated with cognitive decline across patients with dementia and cognitively healthy individuals. This pipeline provides researchers with a powerful tool to study longitudinal brain changes with superior accuracy and sensitivity. It is publicly available and has already been used successfully, proving its utility.

## INTRODUCTION

Magnetic resonance imaging is uniquely capable of capturing brain anatomy in vivo in humans with ever-improving accuracy ^1^. MRI-derived measures of brain morphometry have been demonstrated to be powerful biomarkers capturing differences and changes associated with plasticity ^2,3^, aging and development ^4,5^, and disease and disease progression ^6–8^. Deformation-based morphometry (DBM) is a registration-based technique that uses local volume changes derived from nonlinear deformation fields as a means of detecting local morphological differences/changes ^9–11^. DBM does not require prior segmentation or definition of regions of interest, allowing for the detection of morphological changes in brain regions beyond the common techniques of classifier-driven voxel-based morphometry and cortical thickness analysis.

The technique has been extraordinarily useful in studying rodent models and local volume differences ^12–20^ where a readily accessible toolbox is available ^21^ ^22^. In preclinical models, it has been shown that the volume changes detected by DBM are associated with cellular modifications, such as changing synaptic density across brain networks ^23^. While DBM can be used to compare subjects or groups of subjects cross-sectionally, it is particularly powerful when multiple measures of the same subject are available to model intra-individual time-dependent changes in vivo ^24^. In humans, DBM has been shown to be sensitive and reliable in detecting local disease progression ^6,25,26^, learning-induced plastic brain changes associated with musical training ^27^, and individual brain changes following neuromodulatory treatments ^28,29^.

Classically, DBM uses registration to common template space (e.g. MNI152 ^30^) to derive the necessary deformation fields ^9,10^. Previous work has shown that registration bias and risk of registration error are larger the greater the differences between the template and the individual brain ^31–33^. These are especially relevant in the case of longitudinal studies, where the focus is on changes that occur within each individual brain. To capture these longitudinal individual changes, the deformation fields are best computed using within-subject registration, as even the use of a group-specific template would introduce bias and registration error.

Here we present a publicly available ^34^ two-level DBM technique using state-of-the-art registration tools ^35,36^ designed to capture within-subject changes accurately and allow for the comparison of individual morphological changes across subjects or groups. This two-level DBM technique first calculates the Jacobian determinants for each individual input MRI at the subject level using the within-subject deformation field; then, in a second step, the Jacobian determinants are co-registered to an unbiased common space to be used for statistical analysis. Using artificially induced volume changes and longitudinal images from the OASIS-2 dataset ^37^, we demonstrate the superior sensitivity and accuracy of the two-level DBM pipeline and its lower probability of false positive results when studying longitudinal changes.

## METHODS

### Implementation

The two-level DBM pipeline is implemented as a Python wrapper around antsMultivariateTemplateConstruction2.sh ^38^ from the Advanced Normalization Tools (ANTs) ^39^. In the basic implementation, template construction is applied at two levels: the first level (typically within subject) is computed, generating a subject model and encoding the volumetric differences between input images in a deformation field relative to the subject-specific template (Figure 1). The first-level average templates are then fed into a second-level model build, which constructs an average representation of the population. Two types of jacobian determinants are then computed at the first level, an absolute jacobian, encoding voxel-wise, the nonlinear deformation field, combined with the affine determinant, and a relative jacobian, where residual affine components present in the nonlinear deformation field are removed. The Jacobian maps are then resampled into the final common space and smoothed with a Gaussian kernel with a full-width-half-maximum of twice the smallest voxel size. An optional feature allows the unbiased average to be registered to a common space, such as an MNI model. The Jacobians can then be resampled into that space without contaminating estimated volume differences.

**Figure 1:**
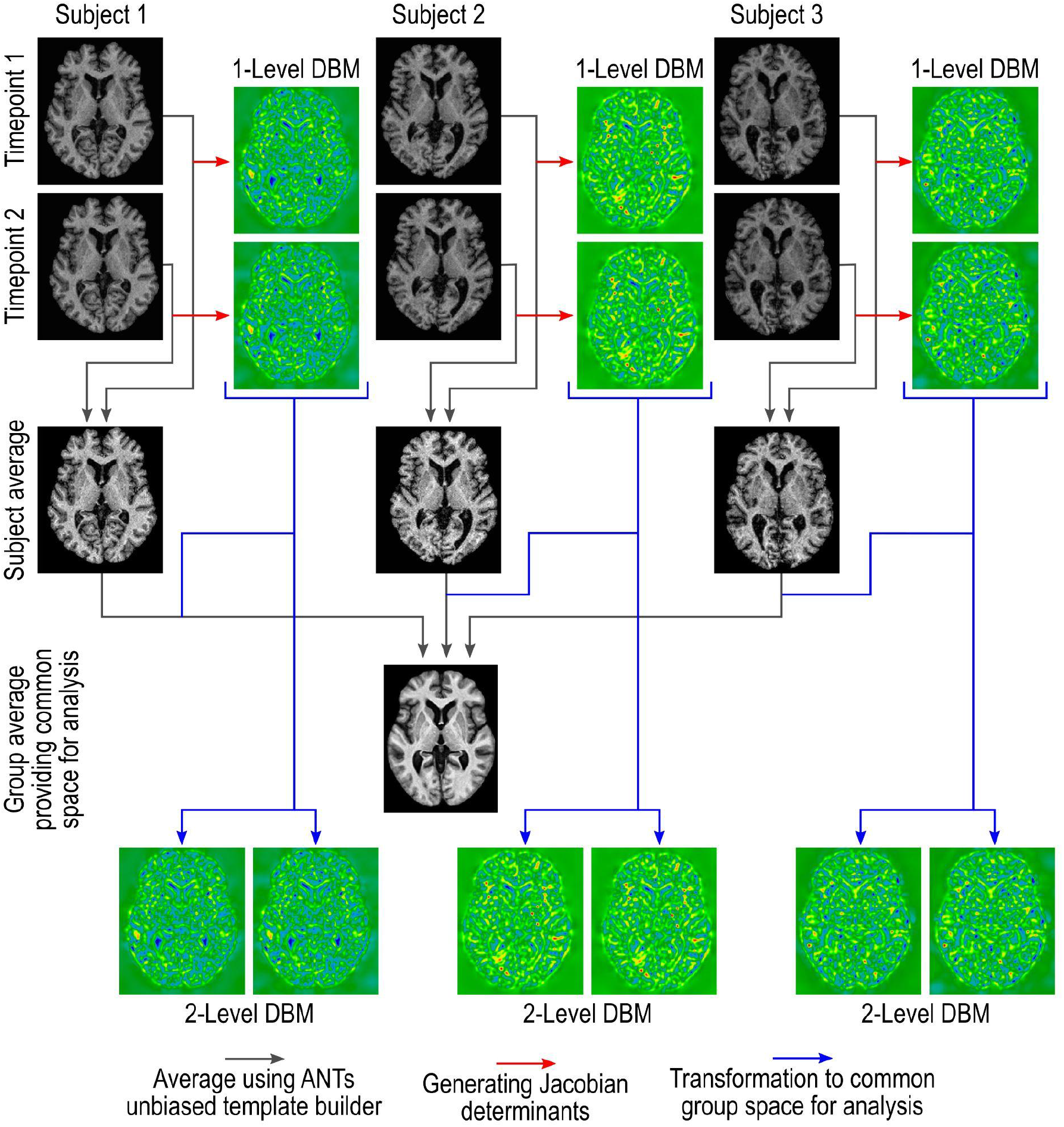
Illustration of the processing underlying the two-level DBM pipeline proposed. Jacobian determinants for each individual input MRI are calculated at the subject level to capture differences and changes with maximal accuracy. The subject-level average brain is then used to generate an unbiased common space, and the Jacobians are transformed into that space for statistical analysis.

### Validation

#### Recovery of artificially induced volume differences

In order to test the sensitivity of this pipeline to volumetric differences in both one-level (i.e., cross-sectional, registration to one common template) and two-level (i.e., longitudinal, within-subject calculation of Jacobian determinants, followed by transformation to common space) implementations, we induced known volumetric changes in the left habenula and right anterior caudate in a set of T1w MRI scans and then used the various DBM pipelines to detect and recover those changes.

20 randomly selected T1w (11 female, age: 61-89 years, mean: 75.45 years ±7.95; 9 male, age: 66-84 years, mean: 72.44 years ±5.63) scans from the OASIS-2 dataset were preprocessed (minc-bpipe-library) and affinely registered to MNI space. The left habenula and right anterior caudate were then semi-automatically segmented using MAGeTbrain ^40^. These two structures were chosen as previous work demonstrated that region characteristics such as size have a small influence on registration sensitivity ^41^. The labels were then used to generate deformation fields for specified Jacobian determinants of 0.5%, 1%, 2.11%, 4.47%, 9.45% and 20% (log spaced 0.5% to 20%) using disptools ^42^. The resulting deformation fields were applied to a random subset of 10 of the subjects to produce a pseudo-second timepoint with an enlarged right habenula and reduced left anterior caudate volume of 0.5%, 1%, 2.11%, 4.47%, 9.45% and 20% (see supplementary video for an example of a modified caudate at 20%). The other 10 subjects were duplicated unchanged. The resulting datasets were constructed as a longitudinal study, with the original brains being used as the first time point, and the 10 modified brains used as the second time point while keeping the remaining 10 unchanged for the second time point (Figure 2). The resulting 6 (levels of local volume change) datasets were processed through the pipeline as both two-level (longitudinal within-subject registration to calculate Jacobian determinant, transformed to the unbiased average space for statistical comparison) and one-level (cross-sectional registration with Jacobian determinants generated relative to unbiased group average).

**Figure 2:**
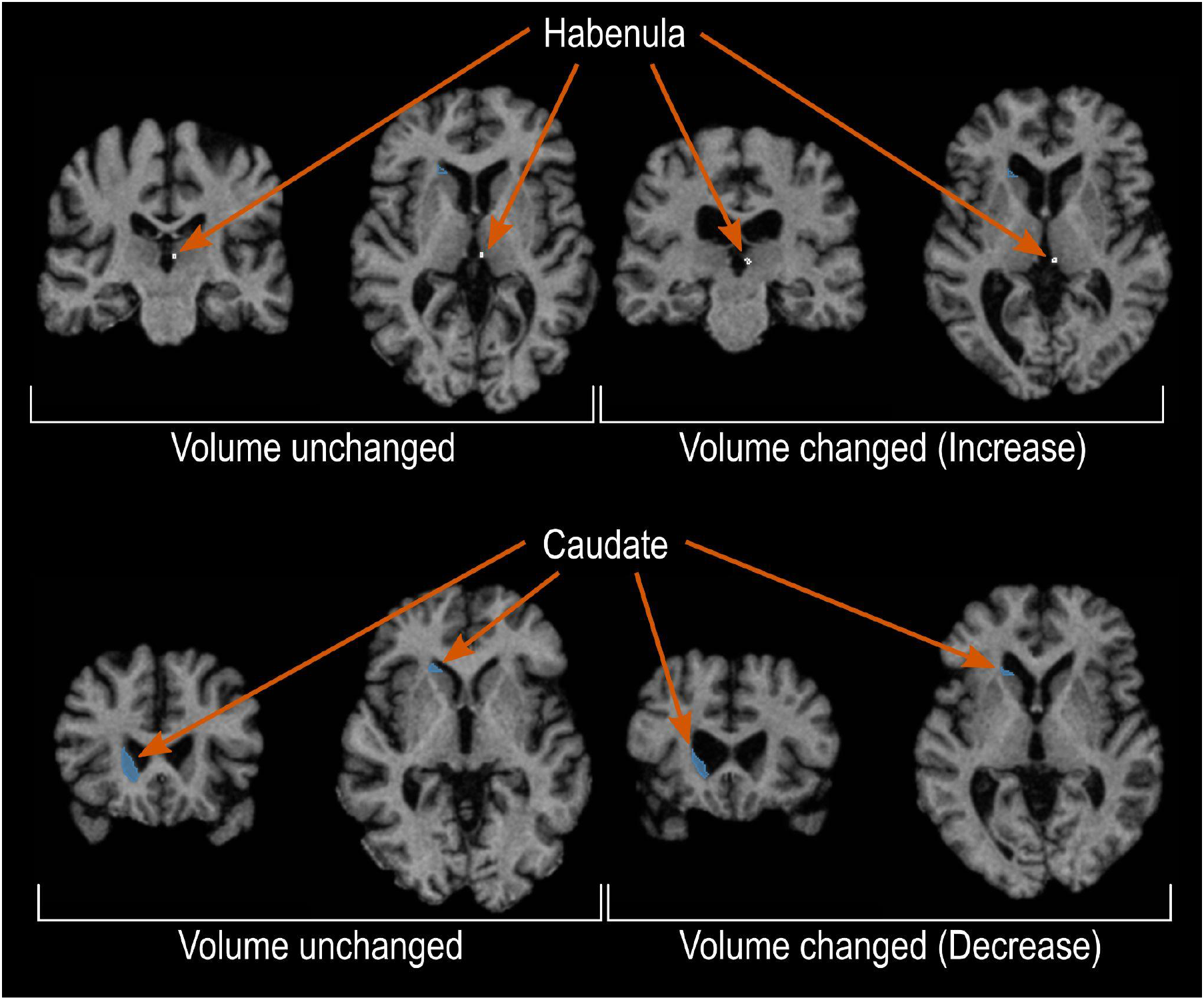
Two sets of brains, 10 brains that remained unchanged and a second set of 10 brains with synthetically induced volume changes to the habenula and caudate regions, were used to assess the sensitivity and accuracy of the two-level DBM. The upper row shows an example of the increased volume in the right habenula region, and the lower row of the decreased volume in the right caudate region.

Voxel-wise statistical modelling was performed using R/3.4.4 and RMINC/1.5.2.2 using mixed-effect linear models, with a fixed effect of (pseudo)timepoint and a random intercept by subject. In addition to voxel-wise modelling, the individual segmentations were resampled into the final template space and merged via majority vote to produce masks. These masks were then used to calculate the volume of the target region by integration of the Jacobian values within the mask to be compared to the true segmentation volumes, thus allowing for an estimation of the effective volume difference captured by each of the pipeline implementations.

### Exploratory analysis of an Alzheimer’s Disease dataset and comparison of DBM implementations

In order to explore the sensitivity of the pipeline in capturing effects in a real dataset, we obtained the OASIS-2 dataset, which contains 150 subjects with 2-5 imaging sessions (mean 2.49, SD 0.69) for a total of 372 longitudinal T1w scans. The dataset consisted of 72 elderly healthy controls (HC; 50 female, Age: 75.5 ± 8.2), 64 participants with dementia (28 female, Age: 75.1 ± 6.7), and 14 patients (10 female, Age: 77.1 ± 7.7) who transitioned during the study between clinical classifications (i.e. from healthy to a diagnosis of dementia). Individuals underwent neurocognitive evaluation at each time point, including the Mini-Mental State Examination (MMSE). The MMSE scores were used for statistical analysis.

The T1W images were pre-processed using minc-bpipe-library (https://github.com/cobralab/minc-bpipe-library), and the skull-stripped brains were processed through 3 variants of the pipeline:

1. The recommended two-level longitudinal pipeline is rigidly initialized with the skull-stripped MNI model and upsampled to 0.5 mm isotropic (mni_icbm152_nlin_sym_09c). In this two-level pipeline, the Jacobian determinants for each individual input MRI are first calculated at the subject level using the within-subject deformation field. In a second step, these Jacobian determinants are then co-registered to an unbiased common space for statistical analysis.
2. The one-level pipeline with the same initialization. Here, the unbiased group average is used as the registration target, and the Jacobian determinants are calculated from the deformation field encompassing the full registration from the subject to the unbiased group average.
3. A modified version of the one-level pipeline to simulate classical DBM, where the previously mentioned model (MNI model upsampled to 0.5 mm isotropic) was used as the full-registration target, and only a single registration was completed.

The third variant aimed at comparing the results to more ‘traditional’ DBM pipelines that use a common brain template ^9,10^.

The 2-level (longitudinal) and one-level (cross-sectional) final averages were post-registered to the MNI152 ICBM09c sym template using antsRegistrationSyN.sh, and transformations were applied to statistical results so that final statistical maps could be compared head-to-head; however, analysis was performed in the unbiased space in each case. Voxel-wise statistical modelling was performed using R/3.4.4 and RMINC/1.5.2.2 using mixed-effect linear models, with a fixed effect of an interaction between days since the first scan with MMSE score, sex, and age at the first scan and a random intercept by subject.

## RESULTS

### Recovery of artificially induced volume differences

Voxel-wise whole-brain mixed-effect models of the artificially induced volume changes are highly sensitive. Statistically significant effects are detected at FDR 5% threshold starting at the artificially induced 2% volume difference. Figure 3A shows a voxel-wise t-statistics map thresholded at 5% FDR. The one-level DBM statistical analysis does not reach statistically significant effects at FDR 5% until the 9% induced volume difference but only detects the caudate difference, not the habenula. Thus from a voxel-wise analysis perspective, the two-level DBM detects volumetric differences that are ∼4.5x smaller compared to a cross-sectional DBM model.

**Figure 3:**
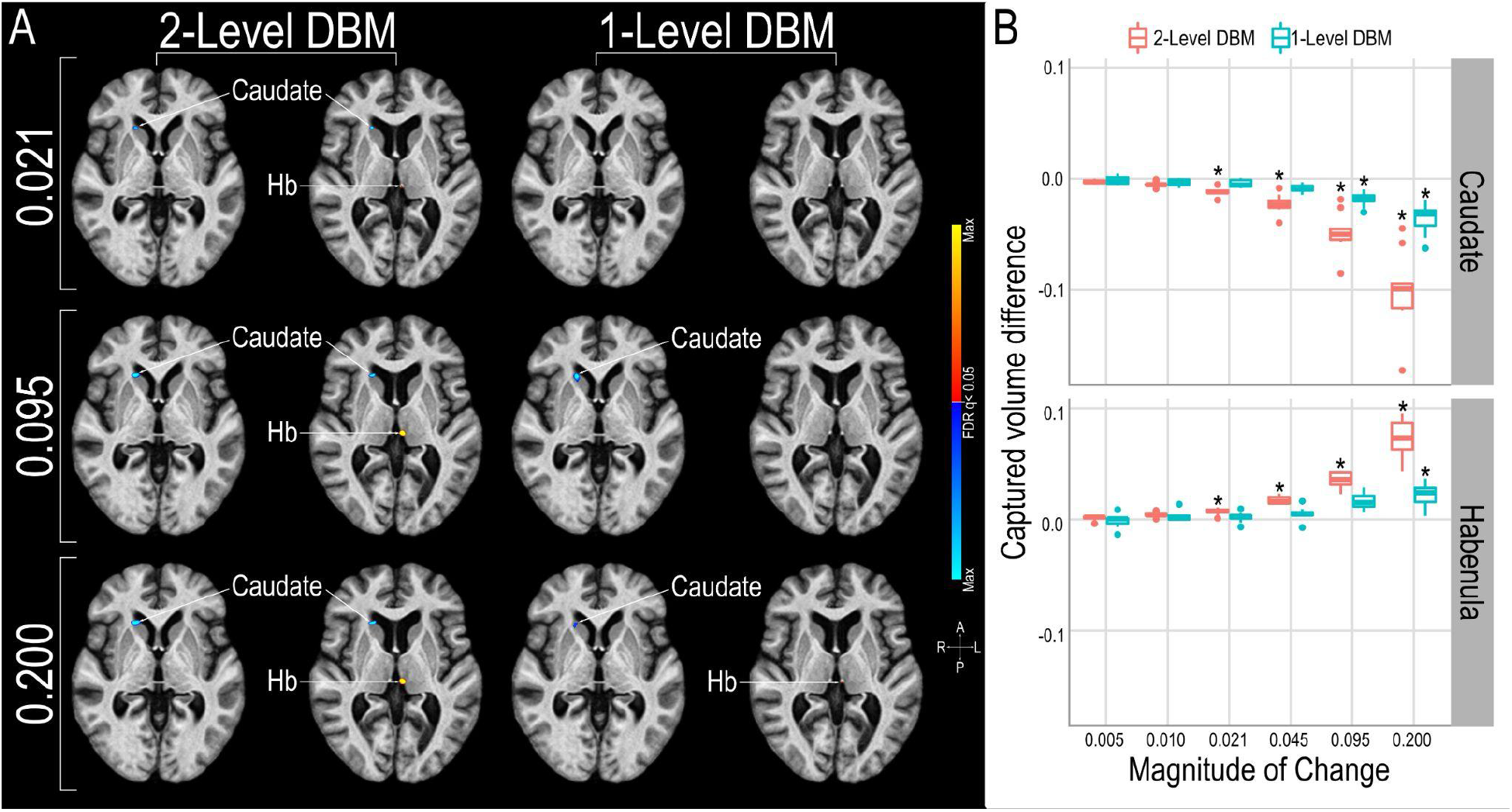
Results of the statistical analysis quantifying demonstrating the superior ability of the two-level DBM to detect and capture the magnitude of the induced volume change. A Sagittal section showing significant (FDR q<0.05) voxel-wise changes. The two-level DBM detects changed volumes at the habenula and caudate regions reliably from a magnitude of 0.021 onwards, while the one-level DBM only detects the induced change at the larger caudate region at 0.095 magnitudes of induced change. The one-level DBM only captures the volume change at the habenula region at an induced change of 0.2 magnitudes (20% increased voxel volume). B Two-level DBM is superior to one level in capturing the magnitude of the induced change at about 50% level.

In addition to examining results on a voxel-wise statistical level, we also examine the inferred volumes estimated through the integration of the absolute Jacobians generated from the DBM within a consensus label. We can examine the resulting volumes with regard to how well they recover the volume difference between the original and induced-volume difference scans. Figure 3B shows the volume change captured by the DBM vs the induced volume difference for the habenula and caudate. The average values are reported in Table 1 showing that the two-level DBM captures the magnitude of the induced change with greater fidelity. Longitudinal volume effect sizes are recovered at approximately 50% of true volume as estimated by the integration of absolute Jacobians within a majority vote mask. Unmodified subjects in longitudinal DBM result in very small volume differences, as expected, whereas in one-level DBM, individual subjects can have volumetric error difference estimates of up to 5%.

**Table 1.** Volume change captured by one and two-level analyses.

| Image type | Brain structure | Magnitude of change | Two Level analysis volume change captured | One Level analysis volume change captured |
| --- | --- | --- | --- | --- |
| Volume Changed | caudate | 0.5 | -0.28 | -0.16 |
| Volume Changed | caudate | 1.0 | -0.50 | -0.33 |
| Volume Changed | caudate | 2.1 | -1.12 | -0.45 |
| Volume Changed | caudate | 4.5 | -2.29 | -0.83 |
| Volume Changed | caudate | 9.5 | -4.85 | -1.84 |
| Volume Changed | caudate | 20.0 | -10.14 | -3.59 |
| Volume Changed | habenula | 0.5 | 0.13 | -0.13 |
| Volume Changed | habenula | 1.0 | 0.42 | 0.26 |
| Volume Changed | habenula | 2.1 | 0.69 | 0.20 |
| Volume Changed | habenula | 4.5 | 1.74 | 0.50 |
| Volume Changed | habenula | 9.5 | 3.58 | 1.63 |
| Volume Changed | habenula | 20.0 | 7.37 | 2.12 |
| Volume Unchanged | caudate | 0.0 (control for 0.5) | 0.01 | 0.04 |
| Volume Unchanged | caudate | 0.0 (control for 1.0) | 0.02 | 0.07 |
| Volume Unchanged | caudate | 0.0 (control for 2.1) | -0.05 | 0.17 |
| Volume Unchanged | caudate | 0.0 (control for 4.5) | 0.01 | -0.34 |
| Volume Unchanged | caudate | 0.0 (control for 9.5) | 0.04 | -0.01 |
| Volume Unchanged | caudate | 0.0 (control for 20.0) | 0.01 | -0.39 |
| Volume Unchanged | habenula | 0.0 (control for 0.5) | -0.02 | 0.06 |
| Volume Unchanged | habenula | 0.0 (control for 1.0) | 0.06 | 0.14 |
| Volume Unchanged | habenula | 0.0 (control for 2.1) | -0.05 | -0.12 |
| Volume Unchanged | habenula | 0.0 (control for 4.5) | 0.03 | -0.24 |
| Volume Unchanged | habenula | 0.0 (control for 9.5) | 0.06 | 0.13 |
| Volume Unchanged | habenula | 0.0 (control for 20.0) | -0.01 | -0.51 |

To examine the possibility of bias in the DBM-based volume estimation, we generated Bland-Altman plots (Supplementary Figure 1) which show the difference between true and estimated volumes compared to the average of those values, bias is indicated by a positive or negative slope. The Bland-Altman plots in both one-level and two-level models show no bias in the estimation of volumes. The range of estimated volumes is substantially larger in the one-level models than the longitudinal models due to scan-wise volume estimates now being contaminated with deformations intended to create the final unbiased average, this is also an excellent visualization of the potential origin of false-positive results in the one-level model.

Considering that the only volumetric differences which exist between the subject-wise scans are known apriori, and the within-subject mixed-effect models, we can examine the potential of false positive findings for the two methodologies. The two ‘timepoints’ are composed of identical images except for the changes to the caudate and habenula region. Computing a main effect of timepoint, we do find sizable groups of significant voxels showing false positive effects using the one-level model (p<0.01, uncorrected). The two-level model, on the other hand, shows only some spurious voxels (p<0.01, uncorrected) (Figure 4). These false positive findings are most likely due to registration errors at the group average stage. As the Jacobian determinants for the two-level model are derived at the first within-subject stage, these errors likely have a much smaller effect on the final statistical comparison.

**Figure 4:**
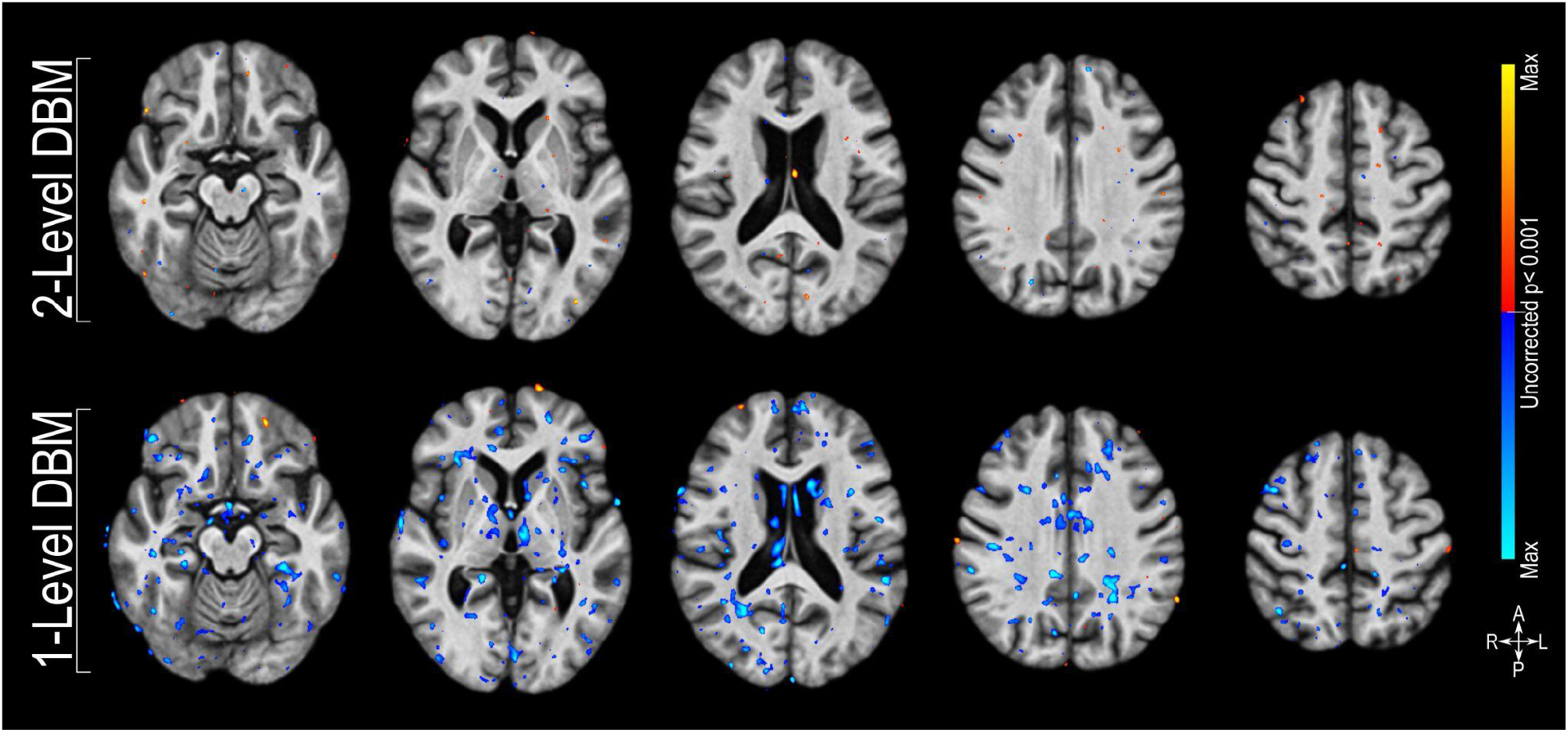
Statistical analysis using Jacobians derived using two-level DBM is less likely to produce false positive findings. Voxel-wise t-statistical maps (p<0.01) of the main effect of time computed using Jacobians derived from the two-level DBM (upper row) and one-level DBM (lower row). Except for the induced volume change to the caudate and habenula region in 10 of the 20 brains, identical brains were used for both ‘timepoints’, no effect of time should be found in this data.

### Exploratory analysis of an Alzheimer’s dataset and comparison of DBM implementations

DBM modelling of the OASIS-2 dataset under the optimized longitudinal configuration reveals a global pattern of relative volume changes associated with a decrease in the MMSE score of participants. As is classically reported, an increase in ventricular volume and CSF space adjacent to the temporal lobes are the most prominent features, indicating an overall reduction in the volume of brain tissue. Secondly, a pattern of gray matter volume reduction in the temporal, frontal and parietal cortices. These effects are highly significant, surviving 1% FDR correction. When comparing the cross-sectional results of the same data, substantially fewer regions of frontal and parietal effects are present at the same threshold, indicating a reduction in sensitivity. Most concerningly, new effects showing an increase in volume of the right superior parietal cortex with increasing MMSE, indicating contamination of the longitudinal brain changes by a group-wise difference present at the start of the study, these effects are false positives, as within-subject effects do not show these findings. Figure 5 shows the head-to-head comparison of the change in MMSE by time interaction in each of the respective unbiased spaces.

**Figure 5:**
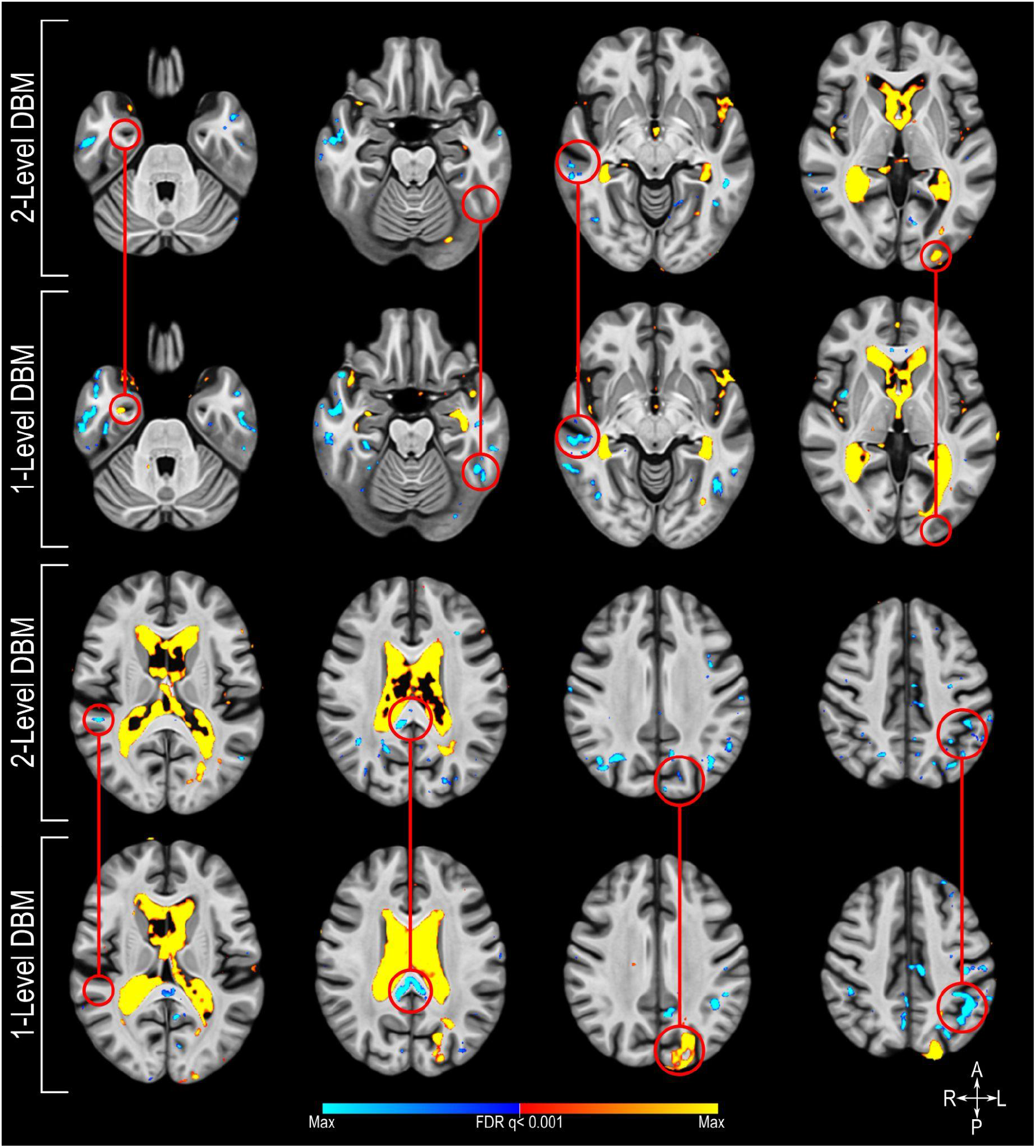
Sagittal sections showing significant (FDR q<0.01) voxel-wise volume changes captured using the Jacobians derived using one and two-level DBM associated with increasing MMSE score over time. Red circles highlight areas where the one-level DBM fails to capture a difference or where it produces spurious results.

To benchmark the sensitivity of the unbiased DBM methods against classical methods, we also derived the Jacobian determinant performing a one-level DBM pipeline direct to MNI space similar to classical DBM methods ^9,10^. The classic DBM using MNI space instead of an unbiased group average shows a pattern of results that is very similar to the unbiased one-level DBM, including many of the same false positive effects (Supplementary Figure 2). However, the sensitivity seems to be much reduced compared to the unbiased one-level DBM, as the sizes of groups of significant voxels are markedly reduced.

## DISCUSSION

The collection of longitudinal data is fast becoming common in both public and private datasets, as well as preclinical, clinical, and population studies. Handling this data in an optimized two-level longitudinal way can yield higher power for a study than using one-level cross-sectional methods. In this study, we introduced a new deformation-based morphometry pipeline which implemented a multi-level unbiased template approach to measuring within-subject whole-brain volumetric change. We validated and compared it against other methods with both synthetic data and a real-world dataset. DBM, as opposed to VBM pipelines, is usable without any tissue classification or atlas priors, allowing for arbitrary contrasts and application to novel species or anatomy. In addition, the absence of a classification stage means that datasets, where successful tissue classification can not be performed, are still usable for DBM. Finally, there are increasing concerns that the classical VBM implementations result in statistical bias due to circularity in their implementations ^43,44^. These prior-free implementations do not rely on such assumptions.

Validation of this pipeline using synthetic data reveals a supremely sensitive tool, approximately 4.5x more sensitive than cross-sectional modelling in detecting volumetric within-subject changes in group-wise comparisons. This result is obtained despite the variability in exact synthetic changes induced by the original manual segmentation process involved in defining the ROI for induced volume change. Very little work has been done in the area of synthetic DBM validation; only van Eede et. al ^41^ examined DBM in a rodent model. There is likely some interpolation error during the creation of the synthetic brains with local volume changes. Therefore, the fact that the two-level DBM implementation is able to capture roughly 50% of the intended change is remarkable.

The two-level DBM not only demonstrates superior sensitivity in detecting longitudinal changes, it also demonstrates a diminished likelihood of providing false positive results. Deriving the Jacobians from within-subject registration allows the minimization of registration bias and risk. Bias and risk grow when using different brain templates for registration, an unbiased group average or the MNI template ^31–33^. This is apparent in the results when examining the much-expanded variance in the volume estimation illustrated in the Bland-Altman plot (Supplementary Figure 1), the false positive findings (Figure 4) and the differences in the pattern of significant voxels in the analysis of the OASIS-2 data (Figure 5, Supplementary Figure 2). These are likely caused by registration errors. The two-level DBM reduces this risk leading to more reliable findings.

The two-level DBM pipeline reported here has already been applied successfully, detecting changes in patients following neuromodulatory treatment for refractory aggressive behaviour ^28^ or mood disorder ^29^, and identifying differences in patients with schizophrenic behaviour ^45^. It has been applied to show how cerebellar and subcortical atrophy contribute to psychiatric symptoms in frontotemporal dementia ^46^. It has also been adapted to investigate brain morphology differences and changes in preclinical models ^47,48^.

## CONCLUSION

Here, we present a two-level DBM technique, where the Jacobian determinants are first calculated at the subject level followed by co-registration to an unbiased common group average. This two-level DBM is particularly suitable for capturing longitudinal intra-individual changes in vivo, being 4.5x more sensitive in detecting longitudinal within-subject volume changes compared to a typical one-level DBM approach and capturing the magnitude of the change much more accurately. The pipeline is publicly available ^49,34^ and will provide users with a superior alternative to analyze brain morphology, especially when using longitudinal data.

## ACKNOWLEDGEMENT

Computations were performed on the Niagara supercomputer at the SciNet HPC Consortium. SciNet is funded by Innovation, Science and Economic Development Canada; the Digital Research Alliance of Canada; the Ontario Research Fund: Research Excellence; and the University of Toronto.

---

**Supplementary Figure 1:**
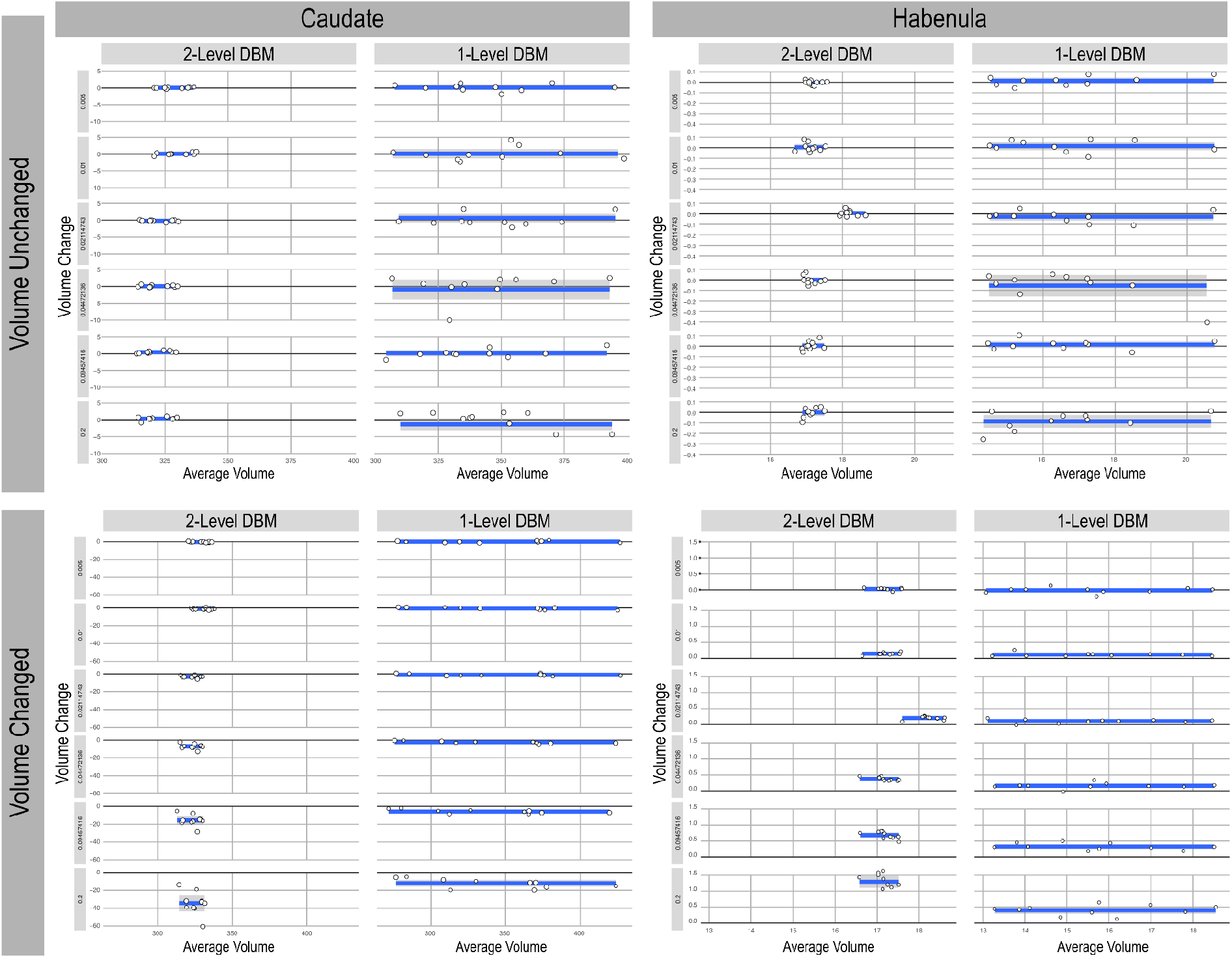
Bland-Altman plots comparing one and two-level DBM across all the levels of induced volume change tested.

**Supplementary Figure 2:**
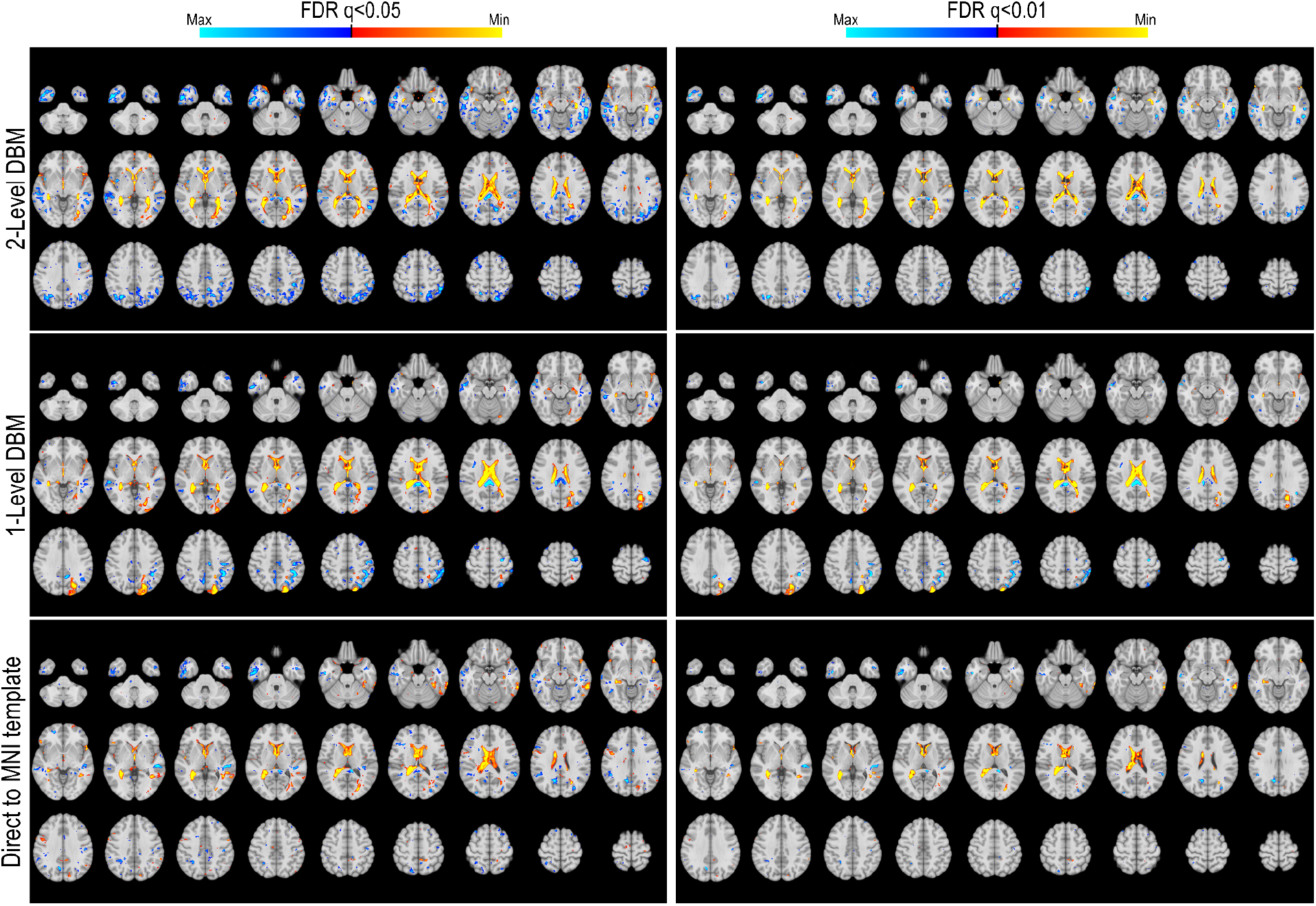
Sagittal sections showing significant (FDR q<0.05 one left, FDR q<0.001 one right) voxel-wise volume changes captured associated with increasing MMSE score over time. The top row reports results using Jacobains generated using the two-level DBM with unbiased averaging - the novel method outlined in this work, the middle row reports results using Jacobains generated using the one-level DBM with unbiased averaging, the lower row reports results using Jacobains generated using the one level DBM to template space - the classical DBM method.

